# Membrane vesicles in freshwater ecosystems: Quantification by flow cytometry and interferometric microscopy

**DOI:** 10.1101/2025.04.10.647775

**Authors:** Viviane Ravet, Violaine Royet, Yasmina Fedala, Claire Hennequin, Didier Debroas, Martine Boccara

## Abstract

Membrane vesicles (MVs) are produced by cells from all domains of life and could be involved in the horizontal gene transfer (HGT). They were found in some environments as seawater and rivers but technological improvements must be required to found out their roles in the ecosystems. In this work, we developed a method using flow cytometry (FC) after lipid staining dye FM4-64 and compared the results to those obtained with Interferometric Microscopy (IM). The abundance of nanoparticles in different lakes determined by the two methods were in agreement. Moreover, nanoparticles with a high refraction index were particularly abundant suggesting that MVs could be associated with several virus particles or to very dense compounds such as iron oxides.

## 1. INTRODUCTION

Viruses are mostly nanoparticles with sizes between 20 and 200 nm and are considered to be the most abundant biological entities on earth (Suttle 2005, 2007). In addition to viruses in the size fraction of less than 0.2 μm, other types of nanoparticle in marine environment, such as MVs have been described (Forterre et al. 2013, Biller et al. 2014). Some Virus like Particles could be in fact MVs. These particles are spherical lipid bilayer compartments with sizes between 20 and 250 nm in diameter generated from cell membranes (Forterre et al. 2013, Biller et al. 2014, Lossouarn et al. 2015, Domingues & Nielsen. 2017). MVs are secreted in the external environment, and are shared by the three domains of life (Gyôrgy et al. 2011, Deatherage & Cookson 2012, Gill & Forterre 2016). MVs are introduced to cargo macromolecules of different compositions (DNA, RNA, proteins, etc.) and are involved in HGT (Domingues & Nielsen 2017). MVs can also carry viruses and act as decoys or Trojan horses (Altan-Bonnet 2016, Tzipilevich et al. 2017). Despite the many ways in which vesicles can affect microbial communities, their abundance and potential functions in environments remain widely unknown certainly because there quantification are biased by the presence of viruses (Schatz & Vardi 2018). By using transmission electron microscopy (TEM) on samples collected from Batata Lake (Brazil), Silva et al. 2016 showed that secretion of outer membrane vesicles is an important cell process of freshwater bacteria. *Prochlorococcus*, the numerically most dominant photosynthetic cell in the oceans releases MVs continually during growth, and these structures were found in abundance in ocean samples (Biller et al. 2014).

In this work, we propose to develop new approaches for counting MVs using fluorescent labelling with lipid dyes and Flow Cytometry (FC), which would be an efficient method to discriminate between various types of biological entities such as viruses and vesicles of the same size. In addition we compared our results with measurements obtained with a new microscope (Boccara et al. 2016). Indeed, recent studies have shown that it is possible to discriminate viruses from vesicles according to their refractive index (Roose-Amsaleg et al. 2017). In addition, we quantified MVs in lakes allowing to give original data since the MVs in these ecosystems are unknown.

## 2. MATERIAL & METHODS

### 2.1 Sampling sites

Four lakes from Auvergne-Rhône-Alpes Region with different trophic status were sampled in 2017: Pavin Lake (45°55’ N and 2°54’ E, oligotrophic, Besse-et-Saint-Anastaise, Puy de dome, France), Aydat Lake (45° 39’ N and 3° 01′ E, eutrophic, Puy de dome, France), Val d’Allier Lake (46° 8’ N and 3° 24’ E, dam lake, Allier, France) and Villerest Lake (45° 58′ 59 N, 4° 02′16 E, dam Lake, Loire, France).

### 2.2 Filtration and concentration of lake samples

Approximately 20 litres of water were taken from the photic zone. Water was successively pre-filtered through nylon membranes (25 μm and 5 μm) to remove suspended organic matter, zooplankton, phytoplankton, protozoa and fungi. The samples were then concentrated using tangential ultrafiltration (Spectrum KROSFlo Minikros Pilot) with a hollow fibre filter (molecular cut-off: 30 kDa, inner diameter: 200 μm, total surface area: 2.6 m^2^) and a transmembrane pressure of 0.07 to 0.1 bars to a final volume of one litre of concentrate (Bisseux et al. 2018). Then, this concentrate was filtrered through a 0.2 μm membrane (Express Plus Membrane polyethersulfone filter, Millipore SA, Saint Quentin, France) and the filtrate was fixed at a final concentration of 1% paraformaldehyde (PFA) and stored at 4°C.

### 2.3 Bacterial culture and vesicle concentration

*Bacillus cereus* was used as a producer of bacterial MVs (Kim et al.2015) to study labelling and FC. Bacteria were grown in culture (500 mL) for 12 hours at 37°C. Two successive centrifugations were carried out on samples, the centrifugation performed at 8000 g and 4°C for 20 minutes to eliminate bacterial cells. The second centrifugation was carried out at 16,000 and 4°C for 30 minutes to eliminate cellular debris.

### 2.4 Fluorescent labelling of membrane and DNA

Samples were incubated for 10 min in the dark at room temperature, at a final FM4-64 concentration of 0.5 μg/ml (Molecular Probes T13320, Eugene, OR). Chloroform treatment was carried out to confirm the lipid composition of the vesicles according to the protocol described by Biller et al 2017. To remove the free DNA fragments labelled with SYBR-Green, the samples were incubated at 37°C with DNase I for 30 min and stopped with EDTA (Invitrogen, Life Technologies, USA). In parallel, to count viruses, samples were diluted with 0.2 μm pre-filtered TE buffer (10 mM Tris-HCL and 1 mM EDTA, pH 8) and stained with SYBR Green I (10,000 fold dilution of commercial stock, Molecular Probes, Eugene, OR, USA). The mix was incubated for 5 min, heated for 10 min at 80°C in the dark and cooled for 5 min prior to analysis (Marie et al. 1999).

### 2.5 Analysis of samples by FC

Particle abundance was determined using a FACS Calibur flow cytometer (BectonDickinson, FranklinLake, NJ, USA) equipped with an air-cooled laser providing 15 mW at 488 nm with the standard filter setup as described by Marie et al. (1999) for SYBR-Green (Marie et al. 1999). Fluorescence intensities were detected by their signature in a side scatter versus green fluorescence plot (530 nm wavelength, fluorescence channel 1 of the instrument) for SYBR-Green and versus red fluorescence plot for FM4-64 (734 nm wavelength, fluorescence channel 3 of the instrument). The flow cytometry list modes were analysed using Cell Quest Pro software (BD Biosciences, version 4.0). Cytometric analyses were also performed on a BD FACS Aria Fusion SORP flow cytometer (BD Biosciences) equipped with a 70 μm nozzle. The laser and filter configuration were as follows: SYBR Green I was excited at 488 nm and fluorescence was collected with a 502 long pass (LP) and a 530/30 band pass (BP).

FM4-64 was excited at 561 nm and fluorescence was collected with 685/50 LP and 710/50 BP. Targeted particles were visualized on a “fluorescence intensity vs. side scatter” dotplot. Data were acquired and processed using FACSDivA 8 software (BD Biosciences). A blank was routinely examined to control for contamination of the equipment and reagents.

### 2.6 Nanoparticle analysis by Interferometric Microscopy (IM)

Five to ten microlitres of each concentrated filtrate was analysed with the interferometric microscope described in Boccara et al. 2016. A stack of 200 images (CMOS camera) was collected. The scattering signals of the diffraction spots produced by the nanoparticles were used to first localize them on each frame. The average number of spots per frame was determined (we estimated the volume of one frame to be 10^−8^ mL) and the concentration of each sample was inferred. The maximum scattering intensity of each particle and its trajectory were computed (Boccara et al. 2016). With these data, we estimated the diameter of each particle from its Brownian diffusion. The scattering signal depended on the refractive index (n) of each nanoparticle (for virus n=1.5) and allowed for discrimination between viruses and membrane vesicles (n<1.5).

Nanoparticle populations were resolved using the Mclust2D package of R software with default parameters (Fraley et al. 2012).

### 2.7 Vesicle and virus imaging by TEM

Fixed freshwater concentrated samples were collected by centrifugation performed at 15,000*g* for 20 min at 14°C directly onto 400-mesh electron microscopy copper grids covered with a carbon-coated Formvar film (Pelanne Instruments, Toulouse, France). Particles were over-contrasted using 2% uranyl salts (w/v; Merck, Darmstadt, Germany). Particles were observed by TEM using a Jeol 1200EX microscope (JEOL, Akishima, Tokyo, Japan) at 80 kV and x 50,000 magnification.

## 3. RESULTS & DISCUSSION

### 3.1 Validation of MVs labelling using bacterial culture

We first analysed the concentrated culture supernatant of *Bacillus cereus* labelled with FM4-64 by FC (Fig. 1). Two populations could be distinguished on the scatter plot: one population (R1 and R2) with a low fluorescence signal and another more complex population (R3) with an intense signal, which corresponded to bacteria. Indeed, both populations disappeared when treated with chloroform (Biller et al. 2017), indicating that R1 and R2 populations corresponded to MVs (Fig. 1). These results suggested that we can identify MVs from environmental samples and count them by FC.

**Figure 1.**
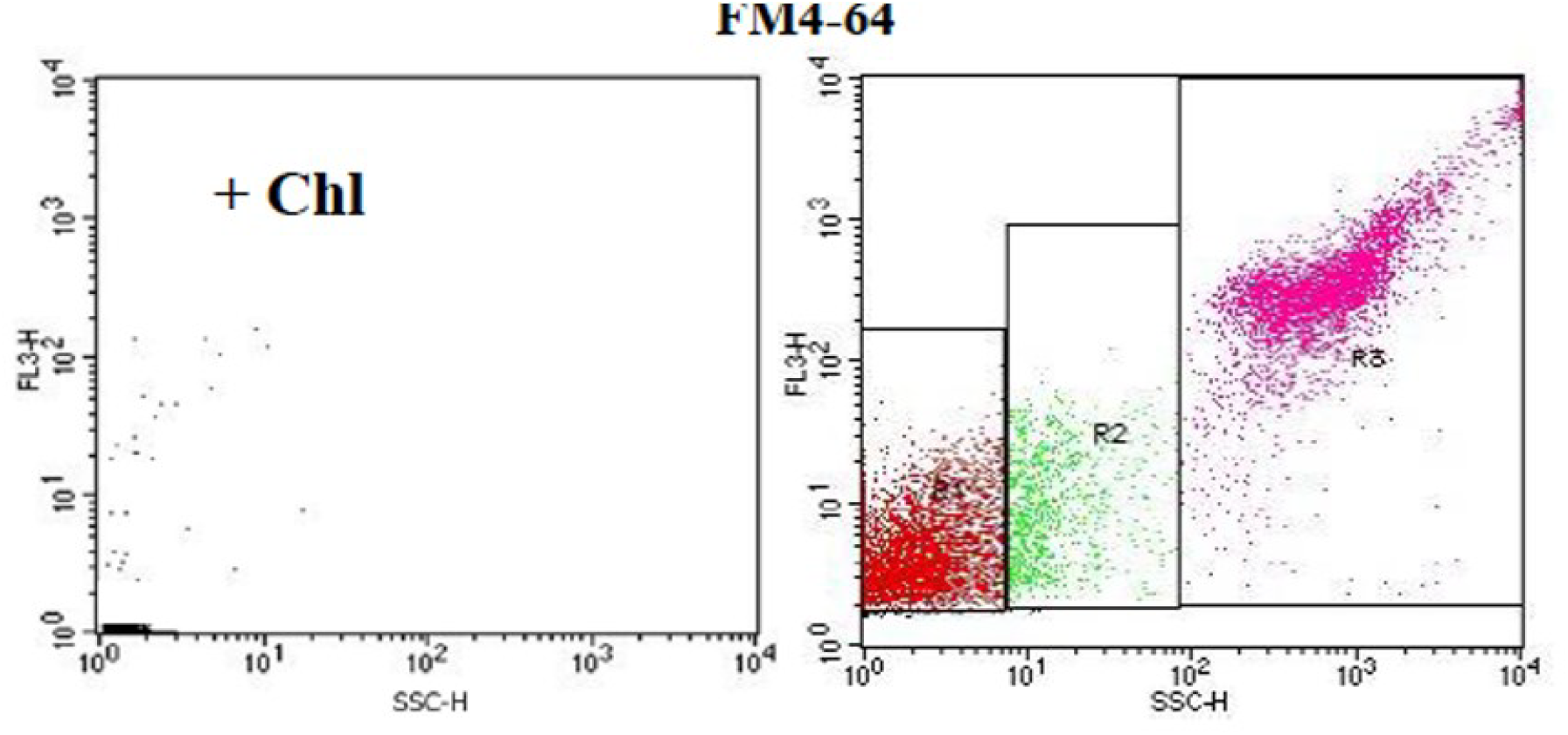
FC analysis of the *Bacillus cereus* culture supernatant: Right panel: Gating of three FM4-64 labelled distinct populations from R1 to R3. Left panel: the same sample was treated with chloroform (+Chl.).

### 3.2 Quantification of nanoparticle populations from lakes by FC

We used FM4-64 and SYBR-Green labelling to analyse sample concentrates (<0.2 μm) from lake ecosystems (Fig. S1). Chloroform treatment was carried out to confirm the lipid composition of the vesicles (Fig. S1). In parallel, the same samples were labelled with SYBR-Green and heated to count the percentage of viruses in the different samples. We observed a decrease in emitted fluorescence when using double labelling (FM4-64 and SYBR-Green) showing likely an interaction between DNA and membrane markers (data not shown). Samples were thus labelled and analysed independently. The results are presented in two separate scatter plots, with the fluorescence intensity as a function of the side scatter (SSC). For example, scatter plots from Aydat Lake are presented in Fig. S1. For quantification, the plots were divided into three regions (R1 to R3) according to their SSC. For nanoparticle quantification, only R1 and R2 of FM4-64 and SYBR-Green labelling (Fig.S1) were taken into account and the results are presented in Fig. 2A (see below).

**Figure 2.**
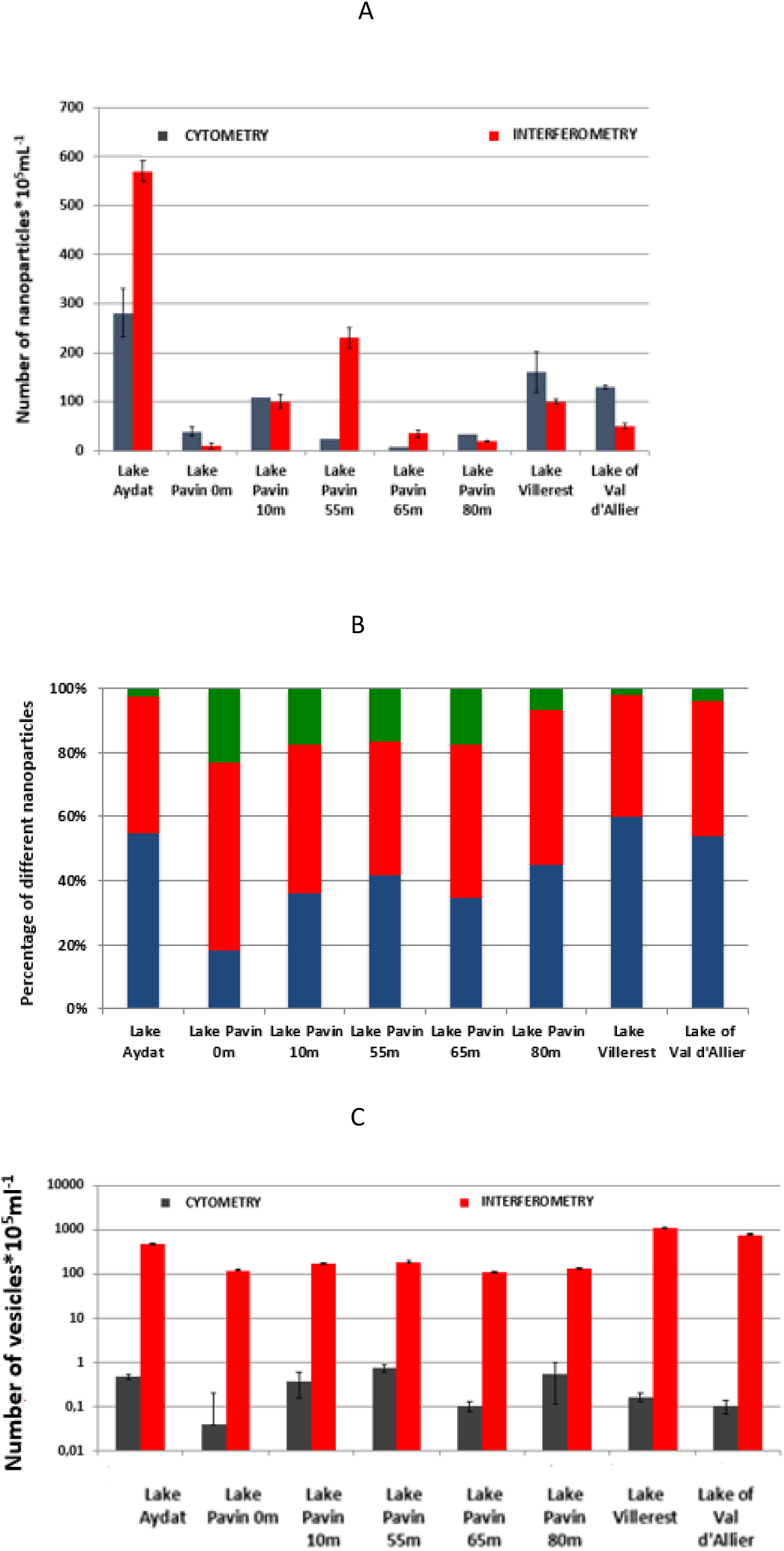
A. Nanoparticle concentrations in the different lakes measured by FC and IM. Number of nanoparticles*10^5^ mL^-1^ in the different lakes estimated by FC (dark blue) and IM (red). The mean values from triplicate and standard deviation are plotted. **B. Histogram of the different classes of nanoparticles determined by IM** The proportions of nanoparticles with different refractive indices were resolved by MClust 2D. Blue: empty vesicles; red: viruses; green: nanoparticles with a high refractive index (1.6 to 2). For Pavin Lake at a depth of 10m, the green fraction corresponded mostly to nanoparticles with a refractive index of approximately 1.6, while nanoparticles with refractive index of 2 represented only 1% of all the particles. **C. Concentrations of MVs measured by FC and IM** The concentrations of MVs were estimated by IM based on the difference in the refractive index (red) and measured by FC (dark blue, FM4-64 labelling) in the various lakes. The mean values from triplicate and standard deviation are plotted.

### 3.3 Quantification of nanoparticles by IM and comparison with FC

The interferometric microscope allowed quantification of the total number of nanoparticles in the sample and analysis of polydisperse suspensions. The abundances of nanoparticles estimated in the same four lake samples were compared using the two method, FC and IM. The concentration of nanoparticles determined by FC was the sum of the number of viruses labelled by SYBR-Green and the number of MVs labelled by FM4-64. The concentrations measured by the two methods were of the same order of magnitude (Fig. 2A). Statistical analysis (ANOVA two ways: lakes and methods) showed that the abundance of the particles varied in relation with ecosystems (p < 10^−8^) but also with the method used (p = 0.003). In addition, there was an interaction between the ecosystem sampled and the method used (IM vs FC). The kind of the particles can explain this result.

IM allowed for the determination of the concentrations of viruses and MVs in a sample. The diameters of the nanoparticles were given by their Brownian displacement (Boccara et al. 2016). In addition, the scattering signal, which depended on the refractive index of each nanoparticle, allowed us to distinguish three types of particles in the lake samples (Fig. 2B): viral particles (refractive index 1.5) approximately 60nm in diameter; other nanoparticles very dispersed in size corresponding to MVs (50 to 250 nm in diameter) with an average refractive index of 1.45; and a third type of nanoparticle that exhibited a higher scattering signal for particles approximately 100 to 200 nm in diameter corresponding to a refractive index greater than to 1.5 (data not shown). The third fraction might have corresponded to aggregates generated during concentration but they contributed to less than 5% of this fraction. We observed a concentrated fraction (<0.2 μm) of Aydat Lake by transmission electron microscopy; that included MVs (blue arrows), viruses (red arrows) and example of two phage particles bound to MVs (Fig. S2). These assemblages could have corresponded to some of the nanoparticles from the high scattering fraction. The high refractive index fraction could also correspond to nanoparticles associated with metal oxides. Indeed, Pavin Lake is rich in magnetotactic bacteria; that produce protein-lipid vesicles; that encapsulate a magnetic crystal made of magnetite (Fe_3_O_4_) or greigite (Fe_3_S_4_) called magnetosomes (Rivas-Lamelo et al. 2017).

In addition, we observed that the numbers of vesicles were significantly different between the two methods and the ecosystems. While the proportion of vesicles in relation to total particles varied between 35 and 65% by the interferometric measurements, it corresponded to only 0.1 to 1% by FC, possibly suggesting that the labelling efficiency was low especially for MVs less than 100 nm in diameter (Fig. 2C).

### 3.4 Conclusions

Few papers describe the abundance of MVs in ecosystems. To visualize the vesicles by FC, we used FM4-64 as a fluorescent membrane marker. We demonstrated that we could label MVs as the label disappeared after chloroform treatment of the bacterial culture supernatant and of vesicles purified Lake samples. FM4-64 labelling showed different populations of vesicles; that differed according to their complexity. We counted the number of MVs that represented a small proportion of the total number of particles.

We used IM (Boccara et al. 2016) to analyse the same lake samples. The abundance of nanoparticles estimated from FC, was different with those estimated using interferometry. Similar results are obtained by Nekrouf et al. (2025) using NTA in mode dispersion or in mode fluorescence to enumerate marine MVs. The abundance of MVs determined by IM was compared with that obtained in other ecosystems, such as in oligotrophic marine environments (Biller et al. 2014) and in the Marne River (Roose-Amsaleg et al. 2017). The interferometry results of the different lakes were of the same order of magnitude as those observed in the Marne River. The estimate of Biller et al. (2014) between 5.10^5^ and 6.10^6^ using nanotracking analysis (NTA) was weaker, but it was a marine environment whose biodiversity was different from the lakes we studied. However, factors of ten to one hundred were observable between the numbers of MVs determined by CF and IM (Fig. 2C). It is possible that the MVs evaluated by FC were underestimate; due to the weak fluorescence intensity with FM4-64 labelling. FC detected only a minority of all cell-derived extracellular vesicles in biological samples measured by NTA (Dragovic et al. 2013). Interferometric microscope, such as NTA, measure vesicle concentrations based on light scatter. We observed differences in the concentration of vesicles (5 to 10 times) between eutrophic lakes and the oligotrophic Pavin Lake using the interferometric microscope, that can differ by their microbial community composition explaining therefore these differences. However, the methods used did not allow discrimination between empty vesicles and DNA containing vesicles, as the latter have scattering signals similar to those produced by viruses.

Interestingly, IM identified nanoparticles with a high refraction index in some samples, which were particularly abundant in Pavin Lake (15% to 20%). Some of these particles could thus be MVs associated with several virus particles (Fig. S2) or to dense compounds such as iron oxides, which is in agreement with the ferruginous nature of Pavin Lake (Rivas-Lamelo et al. 2017).

Here we showed that the two methods of cytometry and interferometry permit estimation of the abundance of nanoparticles in the environment. FC are valuable tools because they can be used to sort particles with different labels. In addition, fluorescence is associated with the interferometric microscope, and with this new set up, we hope to improve the detection of empty vesicles and nucleic acids included in MVs. This last purpose is important to decipher whether MVs can be considered as a fourth way in the horizontal gene transfer as hypothesized by Soler & Forterre (2020).

## 4. Acknowledgments

We thank P^r^ Claude Boccara for helpful discussions and advice, D^r^ Isabelle Mary and Hermine Billard for assistance with flow cytometry, and D^r^ Jonathan Colombet for sample preparation. This work was performed under the auspices of the EC2CO program (from CNRS coordinated by INSU) the VAPOTER project.

## Figure Legends

**Figure S1.**
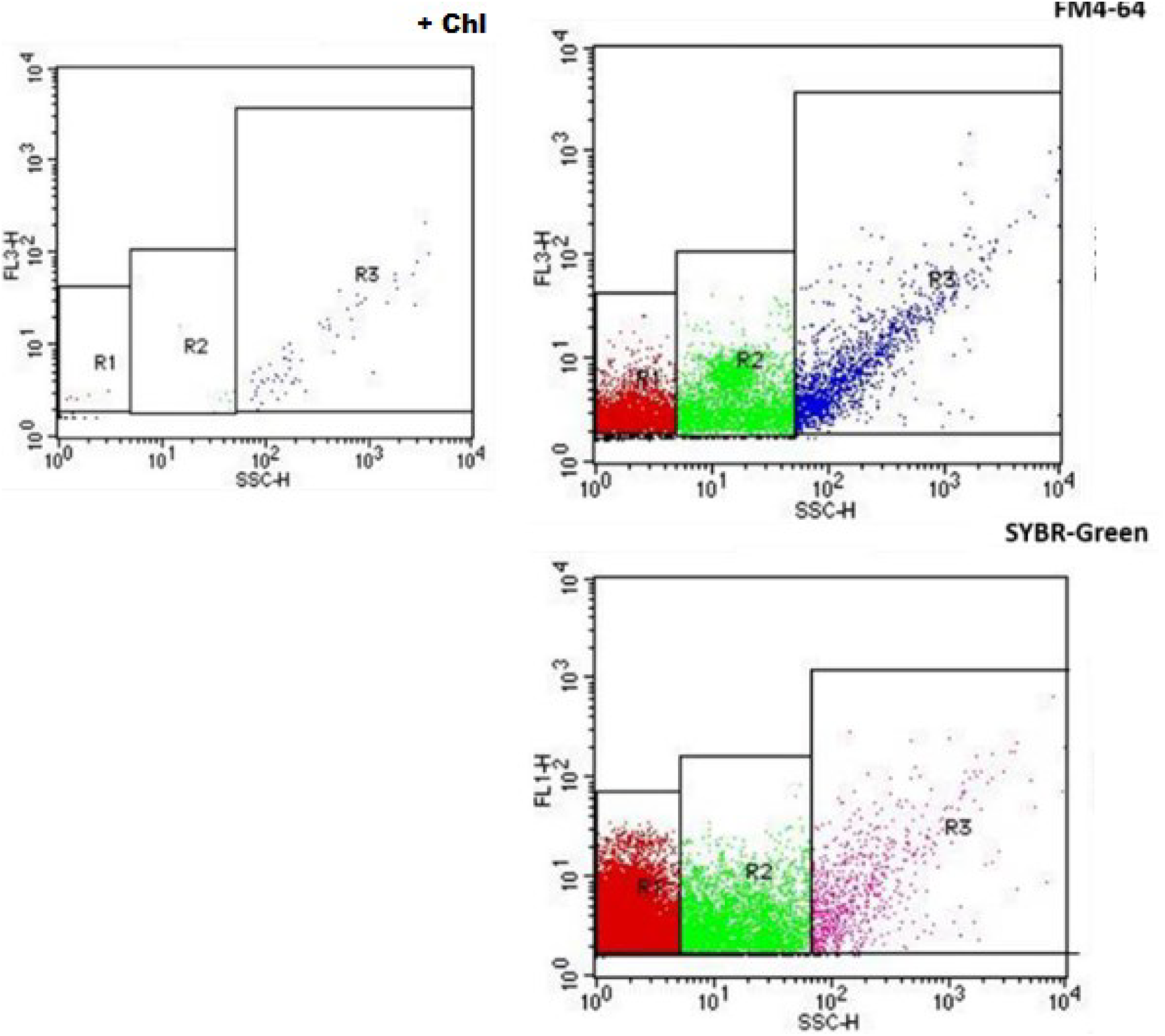
FC analysis of concentrated particles from Aydat Lake. Upper right panel: Gating of three FM4-64 labelled distinct populations from R1 to R3 (upper right panel). Upper left panel: the same sample was treated with chloroform (+Chl.). Bottom panel: Gating of three SYBR-Green labelled distinct populations.

**Figure S2.**
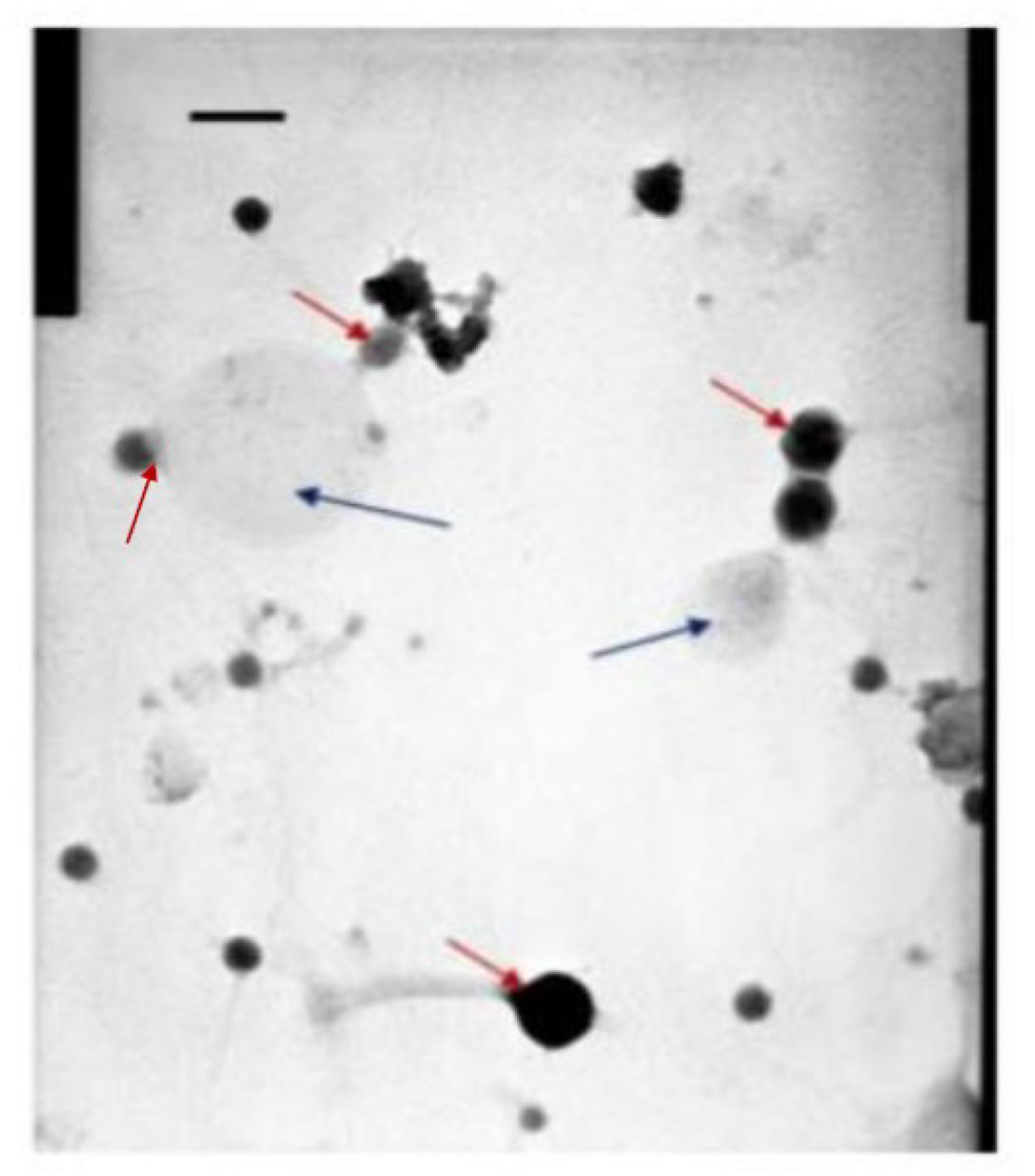
TEM micrograph of nanoparticles from Aydat Lake. The blue arrows show membrane vesicles (MV), and the red arrows show virus particles. Scale bar= 100 nm.

